# Experimental and modeling study of the formation of cell aggregates with differential substrate adhesion

**DOI:** 10.1101/751511

**Authors:** Léo Adenis, Emilie Gontran, Christophe Deroulers, Basile Grammaticos, Marjorie Juchaux, Olivier Seksek, Mathilde Badoual

## Abstract

The study of cell aggregation *in vitro* has a tremendous importance these days. In cancer biology, aggregates and spheroids serve as model systems and are considered as pseudo-tumors that are more realistic than 2D cell cultures. Recently, in the context of brain tumors (gliomas), we developed a new PEG-based hydrogel, with adhesive properties that can be controlled by the addition of poly(L-lysine) (PLL), and a stiffness close to the brain’s. This substrate allows the motion of individual cells and the formation of cell aggregates, and we showed that on a non-adhesive substrate (PEG without PLL is inert for cells), the aggregates are bigger and less numerous than on an adhesive substrate (with PLL).

In this article, we present new experimental results on the follow-up of the formation of aggregates on our hydrogels, from the early stages (individual cells) to the late stages (aggregate compaction), in order to compare or two cell lines (F98 and U87) the aggregation process on the adhesive and non-adhesive substrates.

We first show that a spaceless model of perikinetic aggregation can reproduce the experimental evolution of the number of aggregates, but not of the mean area of the aggregates. We thus develop a minimal off-lattice agent-based model, with a few simple rules reproducing the main processes that are at stack during aggregation. Our spatial model can reproduce very well the experimental temporal evolution of both the number of aggregates and their mean area, on adhesive and non-adhesive soft gels and for the two different cell lines. From the fit of the experimental data, we were able to infer the quantitative values of the speed of motion of each cell line, its rate of proliferation in aggregates and its ability to organize in 3D. We also found qualitative differences between the two cell lines regarding the ability of aggregates to compact. These parameters could be inferred for any cell line, and correlated with clinical properties such as aggressiveness and invasiveness.

## 1 Introduction

The formation of stable aggregates is very common in nature. For example, long-range attraction through chemotaxis can lead to aggregation of Dictyostelium cells [1] or eukaryotic cells during development to form organs and blood vessels [2]). But the formation of aggregates can also arise from Brownian motion and contact adhesion. Numerous examples can be cited, from inert particles such as colloids [3] to living cells, but also in ecology where animals like mussels produce stable patterns by clustering [4]. It has been shown that the living entities can, through this process, optimize at the same time protection against predation and access to food. In cancer, tumor cells circulating in the blood stream form aggregates that will become a metastatic tumor when settling in an organ [5, 6]. The merging of metastatic lumps, forming a larger aggregate, can also occur [7, 8].

It is now recognized that cell cultured in 2D at the bottom of a plastic Petri dish do not behave as they would do in their natural environment. For example, in vitro, an organization in 3D-clusters make the aggregates more resistant to treatments compared to the same cells plated in 2D, in a Petri dish [9]. Several factors can explain this different behavior [10]: first, the fact that the dimensionality is not the same (2D versus 3D) is important; second, cell-plastic interactions are often very strong and prevail over cell-cell interactions; finally, the plastic dish has a very high stiffness, often non realistic (for brain cells for example).

Therefore, new approaches that allow cells to grow in 3D aggregates *in vitro* are being pursued. Aggregates and spheroids *in vitro*, formed on non-adhesive substrates, are considered as pseudo-tumors which can be used to study tumor development in more realistic conditions. Recently, in the context of brain tumors (gliomas), we developed a new PEG-based hydrogel that allows the formation of a tumor-like structure, which can be used to study the effect of drugs in conditions more realistic than those of a 2D Petri dish. In some of those gels, we grafted poly(L-lysine) (PLL), because of its ability to promote unspecific cell adhesion via electrostatic interactions between the polyanionic cell surfaces and the polycationic layer of adsorbed polylysine. The stiffness of the gel can also be tuned, in order to mimic the stiffness of the natural matrix (in our case, the brain) corresponding to a specific cell line. In [9], we chose to study the behavior of two glioblastoma (which is the most aggressive type of gliomas) cell lines. Glioblastomas are currently non-curable and the development of an *in vitro* system that could mimic the development of these tumors could be used in order to test new drugs or radiotherapeutic strategies. Compared to other tissues, the stiffness of the brain is low (lower than 1 kPa [11]), so we chose to design soft gels. We observed a significant difference in cell growth between PLL-containing (adhesive substrate) and PLL-free soft PEG hydrogels (non-adhesive substrate), showing the role of non-specific adhesion factors such as PLL in the migration, proliferation and aggregation in two glioblastoma cell line cultures. More precisely, we showed that on a non-adhesive substrate, the aggregates are larger and less numerous than on an adhesive substrate.

The formation of aggregates has been studied theoretically with a perikinetic equation, first in the context of colloids aggregation [12–15]. In [16], the same concepts are used to study and fit the evolution of the number of cell aggregates on a non adherent substrate. The evolution of the mean projected aggregate area is more difficult to model, especially when aggregates are composed of cells that can deform, contract, modulate their cell-cell adhesion and reorganize in 3D. For example, it has been observed that after there formation, cell aggregates often go through a compaction phase [10, 17, 18] that reduces their projected area.

Other theoretical approaches consider the formation of aggregates under the point of view of phase separation: like two immiscible liquids, when mixed in a liquid medium cells move and seek a lower energy state through adhesion with other cells. The evolution of a system from a state where the concentration of particles is uniform to a final state where patterns appear is a spontaneous phase transition driven by motion of particules, the latter being either passive by diffusion (for example, in colloids), or active (as for mussels or cells) and adhesion. This phase separation corresponding to the formation of aggregates has been described by mean-field models, based on the Cahn-Hilliard equation [19, 20]. In development biology, discrete approaches, in particular with cellular Potts models [21] have been used to model the formation of patterns or the segregation of two cell types in aggregates.

However, to our knowledge, a model that describes the formation of aggregates from the early stages of a population of individual migrating cells to their aggregation and the late aggregate compaction, does not exist. For instance, in [16], the decrease of aggregate area is modeled with an exponential function, but there is no direct connection with processes at the cellular level. In order to be able to describe the individual cells, an agent-based model should be chosen. One advantage of agent-based models is that one can easily implement the local rules of cell-cell interaction ([22–25], for a good review, see [26]).

In [9], we presented the snapshots of already formed aggregates. Here, we add new experimental results by following the whole process, from the early stage where the cell population is composed only of individual cells, to their aggregation, and later to the compaction of the aggregates. We confirm the differential migration and aggregation of cells on the substrates with different adhesivity and for two different cell lines.

We combine these experimental results with a theoretical study based on two models: first, we show that a spaceless model of perikinetic aggregation can reproduce the experimental evolution of the number of aggregates. Second, we developed a minimal off-lattice agent-based model, whose rules are defined in order to reproduce the important phenomena that drive the behavior of cell assemblies: cell and aggregate motion, cell-cell adhesion, cell proliferation and aggregate compaction. We show that this model reproduces very well the experimental temporal evolution of both the number of aggregates and their area, on adhesive and non-adhesive soft gels, for the two cell lines and that it gives access to quantitative values of three parameters.

## 2 Materials and methods

### 2.1 Preparation of the Hydrogels and Glioma Cell Lines

In [9], we showed that the PEG concentration of our artificial substrate optimal for the survival and growth of the two glioma cell lines is around 3% PEG. This concentration corresponds to an elastic modulus around 300 Pa, close to the value measured for brain tissue [27]. We use this concentration in all the following experiments. All the experimental methods can be found in [9]. Briefly: poly(L-lysine) hydrobromide (PLL-HBr 30,000 Da, Sigma-Aldrich, Saint-Quentin Fallavier, France) was first functionalized with an acrylate residue. Hydrogels were prepared from 3% (w/v) PEG-DA 6 kDa precursor (Sigma-Aldrich), dissolved in DPBS with 0.01% (w/v) of DMPA solubilized in VP. Precursor solutions were photopolymerized under UV (UV-LED LC-L1; Hamamatsu, 2 W/cm^2^, *λ* = 365 nm) for 40 s in homemade cylindrical dishes. Photopolymerized hydrogels were then incubated during 1 day in a high volume of DMEM for the hydrogel structure to be hydrated and thermodynamically stable before cell seeding. After this day of hydration and two rinsings with fresh medium, cells were seeded upon hydrogels at 2 10^5^ cells/well (or 10^6^ for the high-density experiments). To avoid cell medium acidification, the cell culture medium was replaced by fresh medium every day.

The two glioma cell lines, F98 from rat model and U87-MG from human glioma, were provided by ATCC (CRL 2397 and HTB-14, respectively). Cells were maintained in Dulbecco’s Modified Eagle’s Medium (DMEM; Life Technologies, Courtabœuf, France) added with phenol red as pH indicator, supplemented with 4.5 g/L of D-glucose and pyruvate, 10% (v/v) fetal bovine serum (Life Technologies), and 1% (v/v) of penicillin and streptomycin antibiotics (Pen-Strep, Life Technologies). Cells were disposed into sterile culture flasks with anti-fungal filters to limit contamination and maintained in culture at 37^*o*^C, under a humidified atmosphere of 95% relative humidity with 5%CO_2_. Cells were replicated when they attained 80% – 90% cell confluence.

In what follows, we call “non adhesive gels” gels composed of PEG only and “adhesive gels” gels containing PEG and PLL.

### 2.2 Microscopy and Image Processing

Microscopy image acquisition was performed using a 10x Nikon water-immersion objective placed on an Eclipse 80i Nikon microscope (Scop Pro, Marolles-en-Hurepoix, France) which was equipped with Differential Interference Contrast (DIC) device. The sample, the stage and the objective were completely enclosed in a chamber that allows the fine control of temperature, humidity and CO_2_ pressure on the living sample (Box and Cube system by Life Imaging Services, Basel, Switzerland). Cells were deposited on the gel surface and rapidly placed under the microscope. We performed time-lapse imaging (1 picture every minute) to follow the formation of the aggregates. Micrographs series were obtained using a Zyla 5.5 MPX Andor SCMOS cooled camera (Scop Pro, France) and MetaMorph acquisition software (Molecular Devices, Sunnyvale, CA).

Image processing (from experiments and simulations) was performed with the Fiji software. Customized Fiji macros were developed to detect cells and aggregates in experiments and simulations. We used Python for further analysis of the data.

### 2.3 The Spaceless Model

The number of aggregates during a perikinetic aggregation process can be predicted by the model of Smoluchowski [12] :

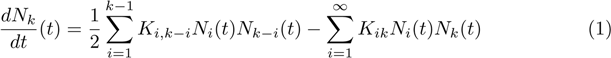

where *N*_*k*_ is the number of aggregates of size *k* and *K*_*ij*_ is the interaction rate between an aggregate of size *i* and one of size *j*. We tried two different scenarii for the aggregation process. In the first scenario, all the aggregates move and interact with other aggregates with the same constant rate *K*. In this case, the model is solvable analytically and the evolution of the total number of aggregates at time *t, N* (*t*), is described by the equation: 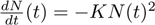, and thus *N* (*t*) = *N*_0_*/*(1 + *K N*_0_ *t*). This model was used in [16] to fit the evolution of aggregate number on a non-adhesive substrate.

In the second scenario, only individual cells move and can interact with other individual cells or with aggregates (*K*_1*j*_ = *K*_*j*1_ = *K*_1*i*_ = *K*_*i*1_ = *K*_1_ ∀*i, j* and *K*_*ij*_ = 0 if *i >* 1 and *j >* 1). The total number of aggregates, *N* (*t*) is given by the following equations, that we solved numerically:

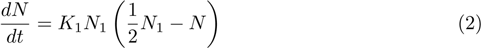

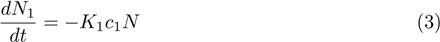

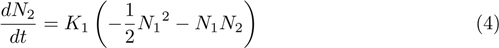

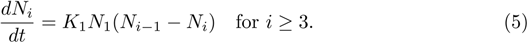

where *N* is the total number of aggregates.

### 2.4 The Spatial Model

We define an agent-based model, that involves a collection of agents evolving in a continuous 2D surface. Each agent is a cell, modeled by a disk. The disk radius is the same for all cells in a given simulation and its value stays constant during the simulation. The simulation space is lattice-free, i.e. the position of each cell is an ordered pair of real numbers corresponding to the coordinates of the center of the sphere.

In experiments, the field of view is a rectangle of size 1280 per 1080 pixels (each pixel is a square of side length 0.658 *µ*m). In the simulations, we choose a square as 2D surface by simplicity. Since the experimental field of view represents a small part of the whole surface of the substrate, it is devoid of boundary effects; therefore, we choose periodic boundary conditions in the simulations (walls or other closed boundaries would induce strong artefacts). The unit of length in simulations is set so that the side of the square has the same length as the length of the region observed in experiments (842.24 *µ*m).

At each iteration, all the cells are updated, one by one and in a random order in order to avoid undesirable correlations. At each iteration, each cell participates in the following processes: motion, both individual and collective, influenced by cell-cell adhesion and aggregate compaction, and proliferation. One iteration corresponds to one minute.

For the sake of clarity, in the following, we call “individual cells” cells that are not part of an aggregate (individual cells have no neighbors). An aggregate is thus an assembly of at least two cells.

### 2.5 Rules of the spatial Model

#### 2.5.1 Cell Motion

At each iteration, individual cells choose a direction uniformly at random and move by a step *a*_0_ in that direction. For a given simulation, this step length is constant during time and is the same for all the cells. Once cells are part of an aggregate, they continue to move, and their motion is the composition of two motions, the motion of each cell inside the aggregate and the motion of the whole aggregate.

The diffusion coefficient of a spherical particle of size *r* in a viscous medium is proportional to 1*/r*; thus, for a 2D aggregate comprising *N* cells, it is proportional to 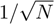. We keep the same dependency here: the motion of the whole aggregate is assumed to be random and the step of this motion is chosen so as to decrease as 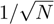.

Regarding the motion of each cells inside the aggregate, the length of the step is chosen as a decreasing function of the number of neighbors *n*: *a* = *a*_0_*/*(1 + *n*^2^). The choice of this function is purely phenomenological. Another choice could have been that of [28] where the probability of migration decreases with the number of neighbors, being proportional to (1 − *q*)^*n*^, where 0 *< q <* 1 is the adhesion parameter and *n* is the number of neighbors. It turns out that the precise choice is irrelevent, since the results do not depend on the exact dependence of the step length on the number of neighbors (for reasonable choices of the corresponding expression).

In the non-adhesive case, individual cells are sometimes subject to a hydrodynamic flux (due to gel local heterogeneity) that can bias the cell motion: when the flux is “on” in the simulation, the direction of the motion in the simulation is limited to a half plane (defined by the direction of the flux), both for individual cells and aggregates.

#### 2.5.2 Superimposition

In order to model the fact that cells are deformable, do contract and can be organized in three dimensions in aggregates, cells are allowed to partially superimpose in the model. The maximum superimposition is quantified by a parameter *α*_max_ that is defined as the ratio of the overlapping length (i.e. the difference between the diameter of a cell and the minimum possible distance between two cell centers) to the diameter of a cell. If *α*_max_ is close to 1, two cells can superimpose almost completely, whereas if *α*_max_ is close to 0, the cells stay well separated. The value of this parameter is constant during a given simulation but can vary from one simulation to an other, in order to model different cell lines. This parameter is similar to the stacking index used in [13] to indicate the formation of aggregates with some vertical stacking, forming multilayered clumps.

#### 2.5.3 Adhesion

If during its motion, the position of a moving cell would break the superimposition rule, the motion is prematurely stopped and the two cells adhere to each other. There is no break-up mechanism and so cells cannot detach from their neighbors (except if its step leads it to a position where it has at least the same number of neighbors). Therefore, if the cell chooses a direction of motion that would lead to a smaller number of neighbors after performing the step, this step is canceled and the cell does not move.

#### 2.5.4 Compaction

We do not have precise details on what happens inside the aggregates during compaction, so we chose to model the effect of cell compaction and 3D organization with a simple yet efficient empirical rule: we bias the individual cell motion in an aggregate towards the center of mass of the aggregate. Since this reorganization is much more visible for larger aggregates, we decided to modify accordingly the cell motion. When the aggregate is very small, its cells may move towards any direction. However, when the aggregate becomes more massive, the cell motion is biased towards the center of mass of the ensemble. At the limit of a very massive aggregate, the motion is possible only in a ± 90*°* sector around the line joining the cell with the center of mass. This bias in the direction is similar to the one in [29] where the direction choice is biased towards the direction with the higher number of cells within a distance corresponding to several cell diameters. The precise implementation of the motion bias does not concern us here: the overall effect does not depend crucially on the details.

#### 2.5.5 Proliferation

We model the cell division by the following process: after the cell has moved it has a certain probability to try to proliferate. That probability per iteration is later on referred as “proliferation rate”. If the cell division process is engaged, a random position around the dividing cell is chosen. If that position is compatible with the superimposition rule (meaning that the distance between the daughter cell and all the other cells should be larger than the minimum distance allowed by the superimposition coefficient), the daughter cell is created. If not, the division process is aborted.

### 2.6 Typical Workflow

Simulations yield the positions of the cells, that are used to produce images representing the same observed area as in the experiment. The size of individual cells in simulations is chosen equal to the mean size of cells in the experiments, so that the simulation results can be analyzed the same way as the experimental results, and we can then directly compare the dynamics of the mean area and of the numbers of aggregates in the two approaches.

## 3 Results

### 3.1 Experimental Observations

On Fig 1 top and on the different movies of aggregation, we observed several phases in the aggregation process (S1 Video for F98 and S2 Video for U87).

**Fig 1.**
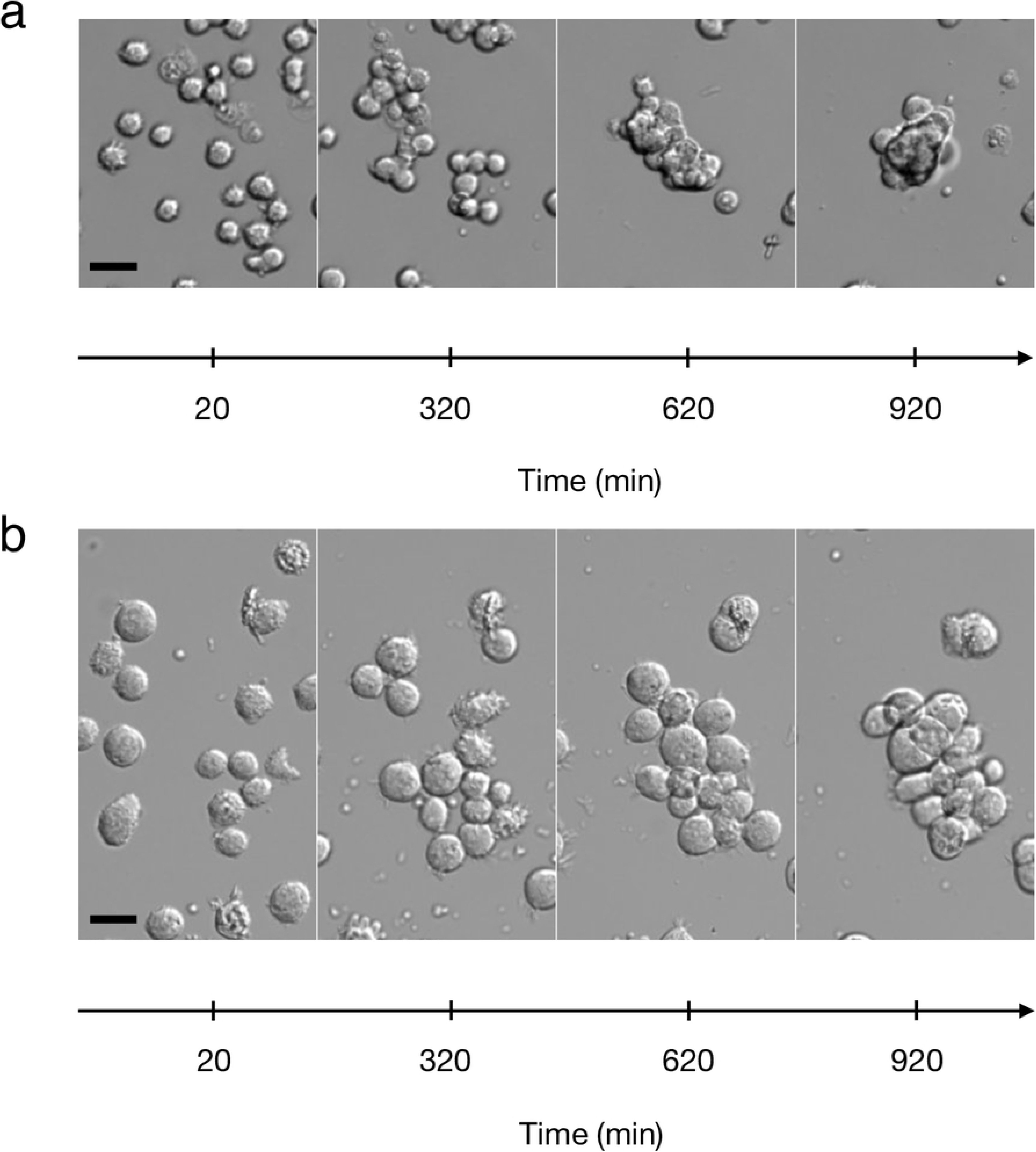
Aggregation process: Images at different times, during the process of aggregation, for the F98 cell line (a) and the U87 cell line (b). The scale bars represent 20 *µ*m.

The first phase is the random motion of individual cells. Cells remain round, do not polarize, and do not adhere strongly to the substrate. However, they move, extend small filopodia, and perform a random motion (see S1 Video and S2 Video). After analyzing the motion of 25 cells (F98) during one hour, we plotted the mean square distance covered versus time and we deduced that the cells have a diffusion coefficient of 1.3 ± 0.27*µ*m^2^ min^−1^ on the adhesive substrate (data not shown), close to the value found in [30].

The second phase is the formation of aggregates: during this random motion, individual cells encounter other cells or already formed aggregates and new aggregates begin to form. When this happens, the individual cell sticks to the other cell or to the aggregate; becoming unstuck is so rare that we neglected this phenomenon.

The third phase corresponds to the dynamics of formed aggregates: they move as a whole, with small aggregates exhibiting a global motion larger than the bigger ones. Moreover, the inside of the aggregates is also dynamic: the cells inside move and reorganize constantly (see S1 Video and S2 Video). During the aggregation phase, which lasts around 2 hours, a few events of proliferation are visible among individual cells. This proliferation continues within aggregates, increasing their size continuously [9].

The last phase corresponds to the compaction of the already formed aggregates: a few hours after the formation of aggregates, they compact and reorganize into a three-dimensional shape. The projection of this shape in 2D is close to a disk. Experimentally, compaction occurs only for the F98 cell line. For the U87 cell line, this cell contraction does not occur and cell aggregates stay in a 2D configuration, see Fig 1.

We measured, in the experimental field of view, the mean area of the aggregates and their number as a function of time, for adhesive and non-adhesive substrates, for the F98 and the U87 cell lines. We define the normalized number of aggregates as the raw number of aggregates divided by the number of individual cells at initial time. In Fig 2, the normalized number of aggregates (top) and the mean area of aggregates (bottom) are represented as a function of time, for non-adhesive (blue curves) and adhesive (brown curves) substrates, for the F98 cell line. The number of aggregates (respectively mean aggregate area) decreases (resp. increases) faster in the case of the non-adhesive substrate.

**Fig 2.**
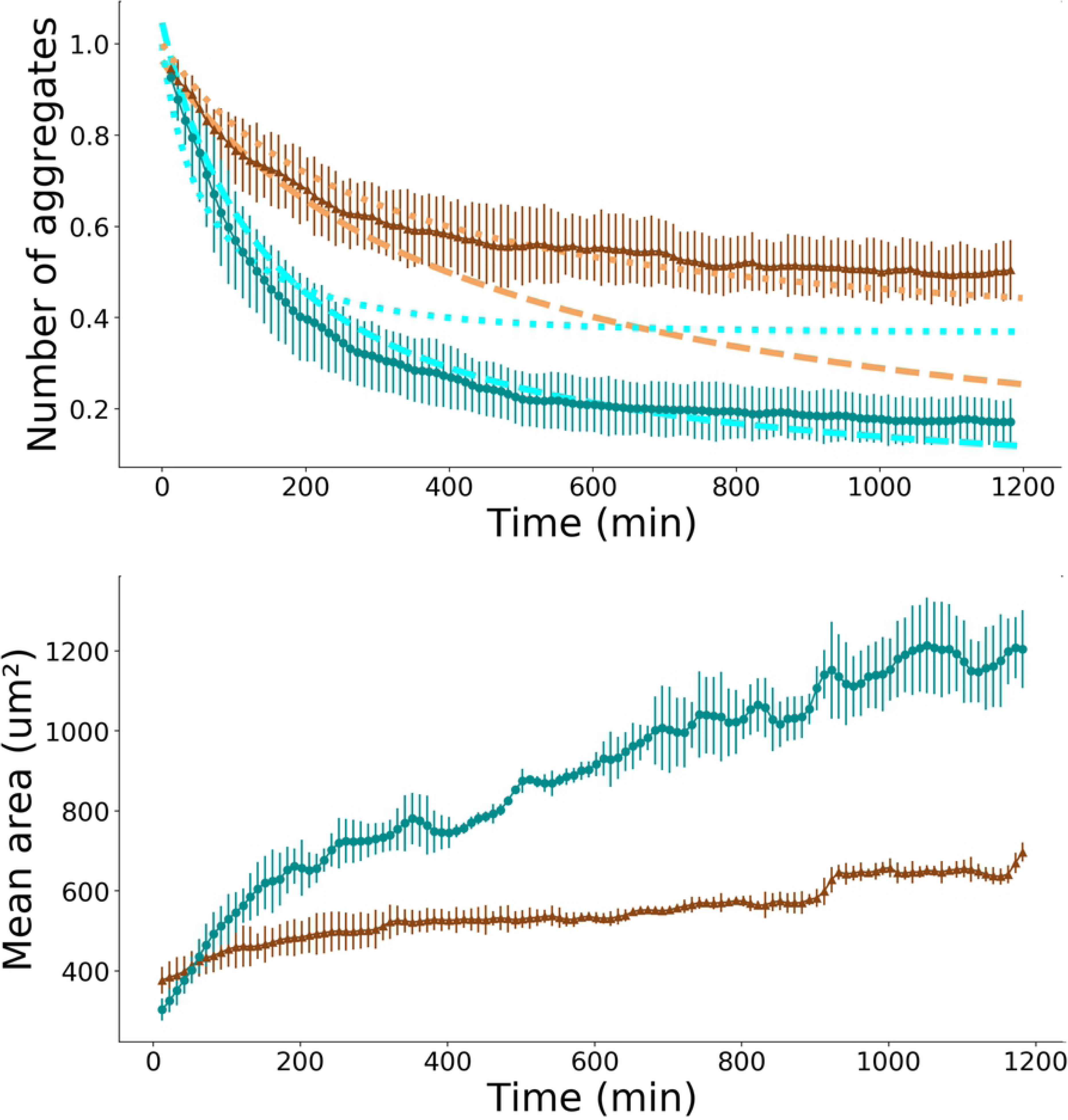
Temporal evolution of the number and area of aggregates. Normalized number of aggregates (top) and mean aggregate area (bottom) as a function of time, for the F98 cell line, in the case of adhesive substrate (brown triangles) and non-adhesive substrate (blue circles). The mean value is calculated from three experiments in each condition, error bars represent the standard deviation. The brown and the cyan dotted lines correspond to the best fit of the experimental data with the numerical solution of the spaceless model based on Smoluchowski equations in the case where only individual cells move. The brown and the cyan dashed lines correspond to the best fit of the experimental data with numerical solution of the spaceless model based on Smoluchowski equations in the cas where the aggregation kernel is a constant.

### 3.2 Comparison Between Experimental Data and the Spaceless Model

We compared our experimental results with the theoretical model for aggregation developed by Smoluchowski [12]. In the case of a non-adhesive substrate, we found that the best fit was obtained with a constant aggregation kernel *K*. From the fitting procedure, we found *K* = 2.6 10^−13^ m^2^s^−1^, see Fig 1, top (the dashed blue curve is obviously a better fit of the experimental data than the dotted blue curve).

In the case of an adhesive substrate, we found that the best fit is obtained with the solution of the equations corresponding to the scenario where only individual cells can move and interact with the other aggregates. In this case, we found *K* = 6.4 10^−14^ m^2^s^−1^=3,8 *µ*m^2^min^−1^, see Fig 2, top (the dotted brown curve is obviously a better fit of the experimental data than the dashed brown curve). This value is of the same order of magnitude as the cell’s diffusion coefficient on adhesive substrate calculated above.

### 3.3 Comparison between Experimental Data and the Spatial Model

We model two cell lines: F98 cells (13.2 *µ*m mean diameter) and U87 cells (21.1 *µ*m mean diameter).

#### 3.3.1 Qualitative comparison

The rules of our model (sketched in Fig 3 (a) have been defined in order to mimic what happens in the experiments: therefore, in the model, cells can move, adhere to other cells, form aggregates that can contract subsequently, and proliferate.

**Fig 3.**
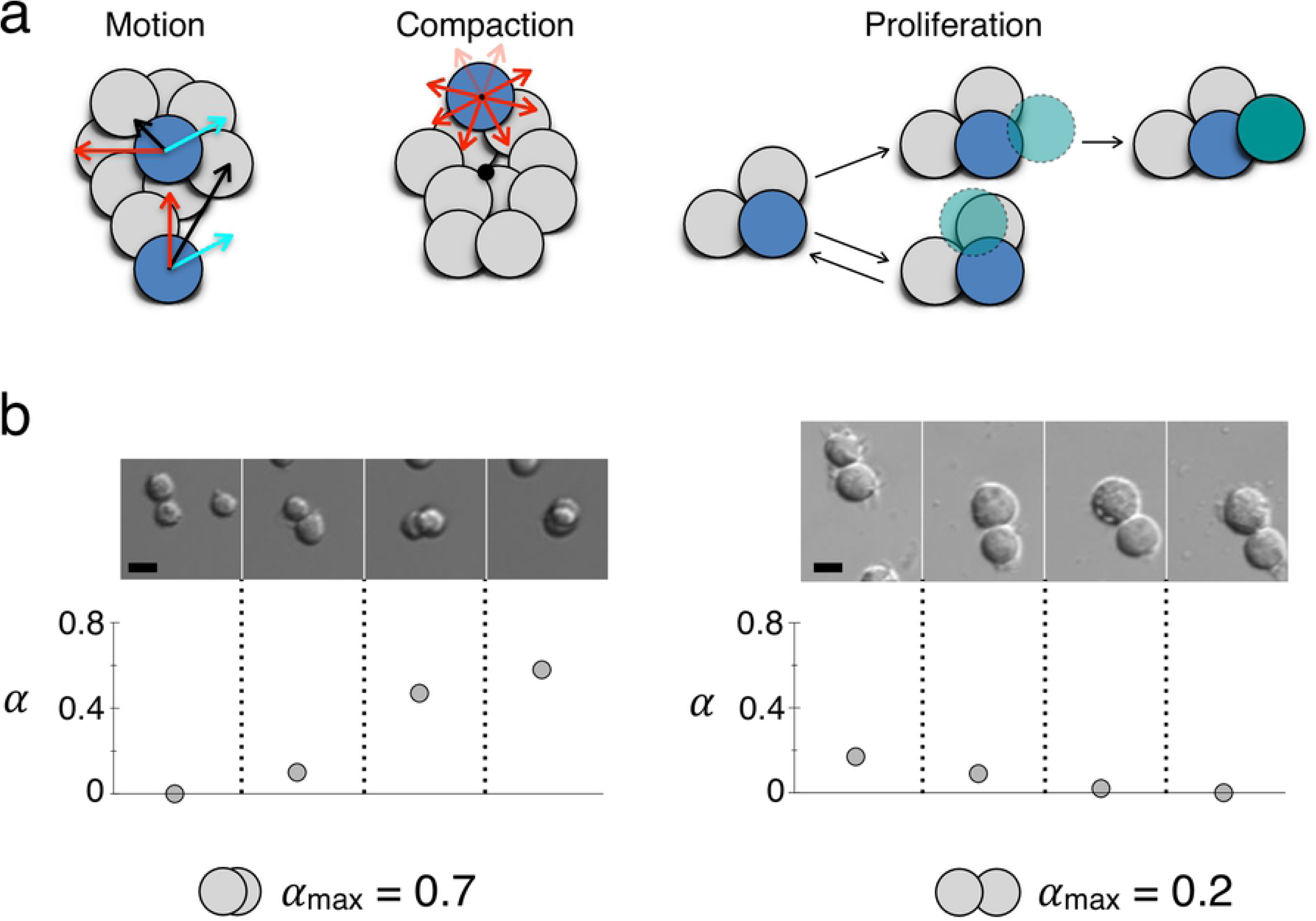
Rules of the spatial model. (a) Left: cell motion (black arrow) has two components, the first one (cyan arrow) is common to all cells forming an aggregate and is decreasing as the aggregate is growing, the second one (red arrow) is the individual motion of each cell, whose length becomes smaller as the number of neighbors increases. Center: the rule for the aggregate compaction stipulates that the bigger the aggregate the more biased towards the center of mass of the aggregate (the black circle) the individual cells’ motion is. Right: proliferation; in the upper sketch, the foreseen daughter cell (the green cell with a dashed border) is really created, whereas in the lower sketch, it is too close to other cells so the daughter cell is not created. (b) Top: Images of two-cell aggregates, for the F98 cell line (left) and U87 cell line (right), and the values of the corresponding superimposition coefficient *α*. Bottom: schematic representations of the two values *α*_max_ that were chosen for the F98 cells (*α*_max_ = 0.7) and for the U87 cells (*α*_max_ = 0.2). The scale bars represent 10 *µ*m.

In S1 Video, it is clear that cells move inside an aggregate, and since aggregates also have a motion on their own, the motion of each cell should be a composition of the two motions, see Fig 3 (a), left.

It is well known that aggregation limited by diffusion leads to clusters with fractal shape [14, 15, 31]. In our case the number of cells is not large enough to lead to fractals, but without any rule of contraction, especially in simulations with a high initial cell density, aggregates have the shape of long and branched thick filaments and do not organize themselves into more compact shapes. But in experiments at high cell densities, after about 12 hours after the beginning of the aggregation process, the aggregates compact and become more circular (in 2D) (see Fig 4 (a), middle). The effect of the compaction rule in the case of a high initial cell density is visible on Fig 4 and leads to aggregates that have a compact shape, close to the experiments.

**Fig 4.**
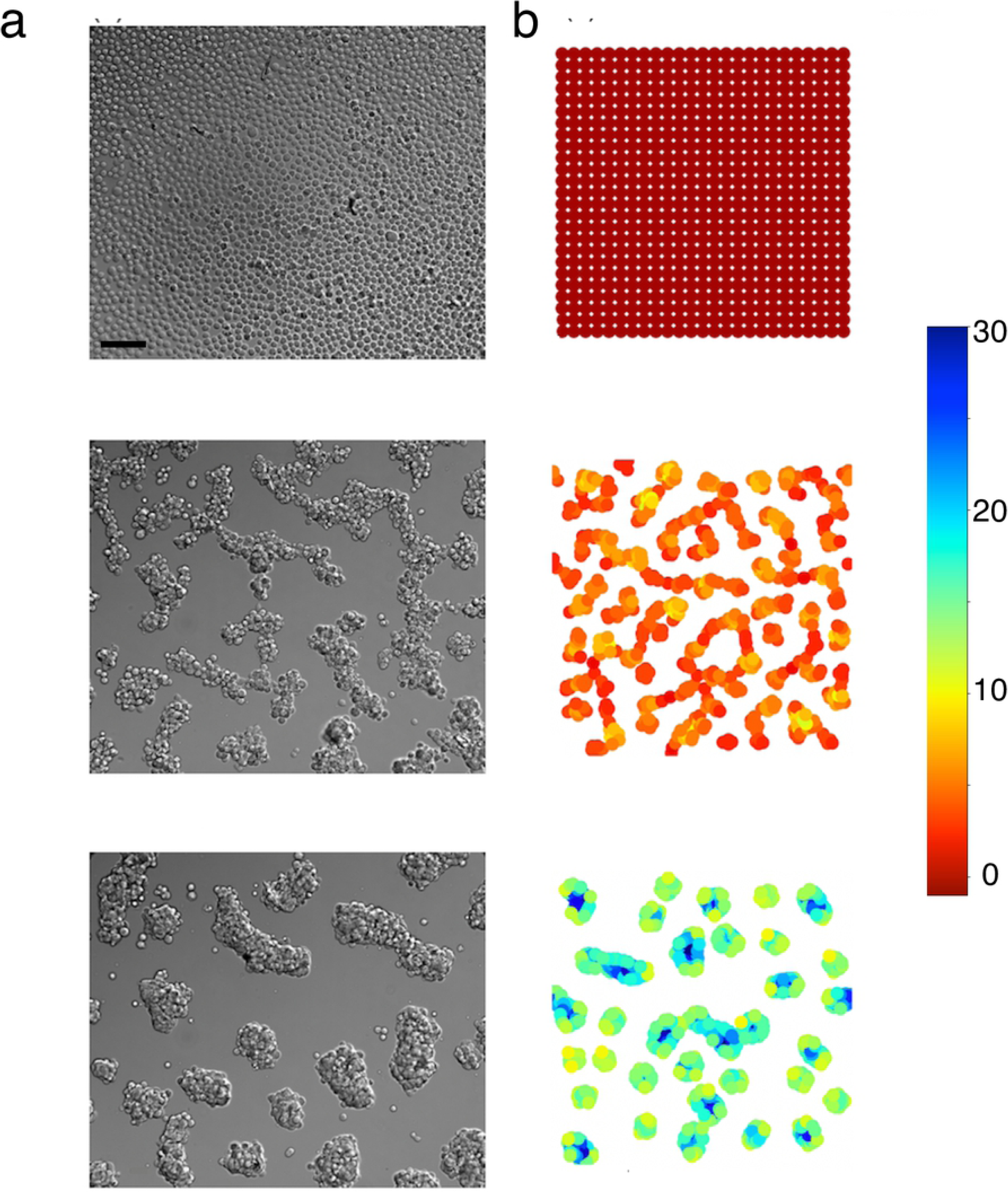
Compaction at high cell density: (a) experiments, (b) simulations. Top: *t* = 0; middle: *t* = 6 h; bottom: *t* = 12 h. The black scale bar on the bottom of the image at the top, left, represents 50 *µ*m. Cell color in simulations is a function of the number of neighbors (the scale is on the right).

The qualitative comparison of the experimental and the simulation results can be performed from Fig 5, which shows side by side an experiment and a typical simulation result, obtained with our model.

**Fig 5.**
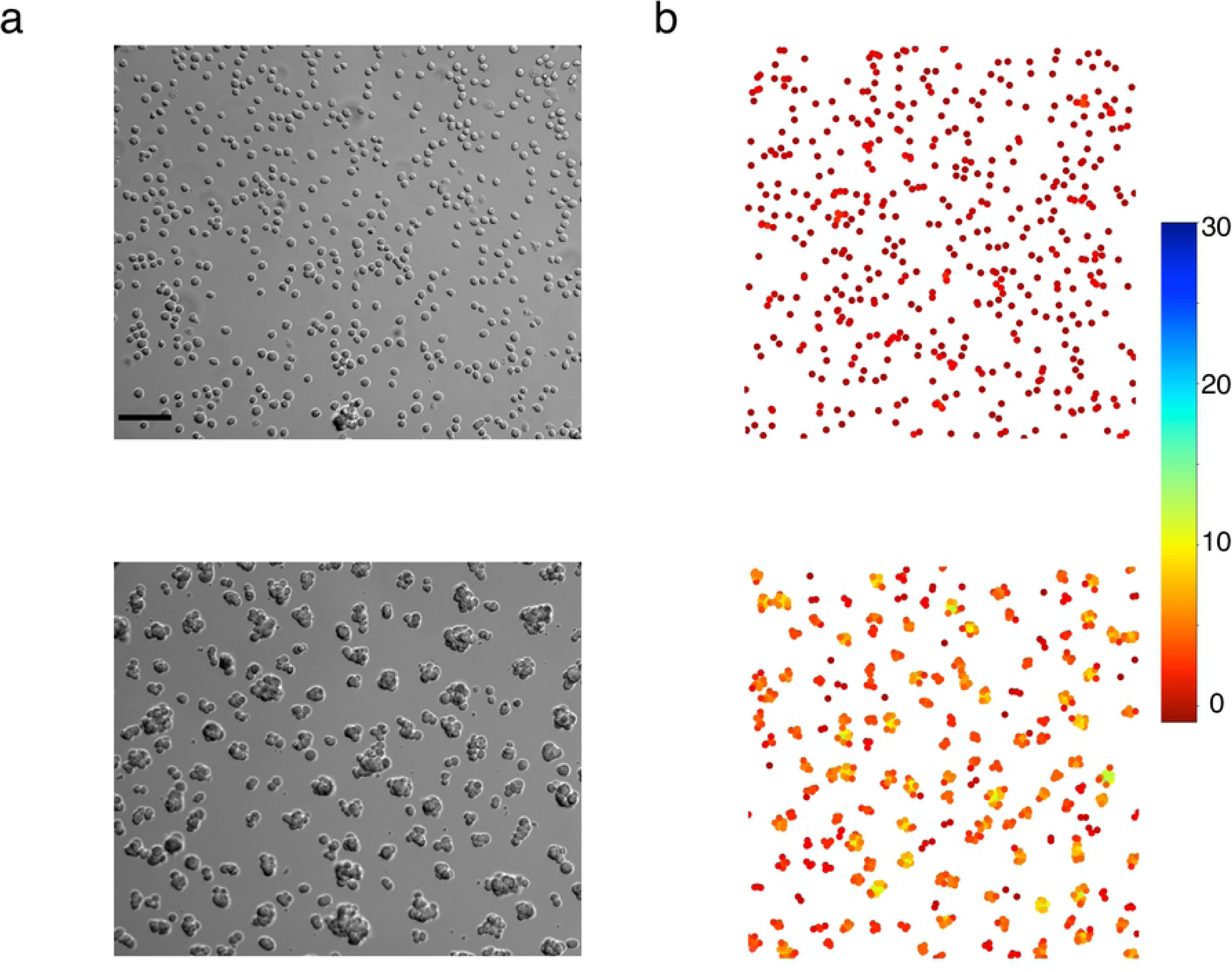
Qualitative comparison between experiments and simulations: (a) experiment and (b) corresponding simulation; top: initial states of an experiment and a simulation respectively; bottom: final state of the same experiment and simulation (12h later, or 720 iterations with our time calibration). The black scale bar on the bottom of the top left image, represents 50 *µ*m. Cell color in simulations is a function of the number of neighbors (the scale is on the right).

### 3.4 Quantitative Comparison between Experimental Data and the Results of the Spatial Model

The experimental results consist of 9 24-hour experiments: 6 experiments for the F98 cell line (3 in the adhesive and 3 in the non-adhesive condition), and 3 experiments for the U87 cell line (one in the adhesive and two in the non-adhesive condition). We decided to fit each experimental curve (not only the mean), so the parameters were independently set for each experiment.

#### 3.4.1 Choice of Parameters

The number of initial cells *N*_0_ was deduced from the first image taken in each 24-hour experiment and was used as the initial condition of the corresponding simulation, a uniformly random distribution of *N*_0_ individual cells. The superimposition parameter was chosen constant for all the experiments with the same cell line. There is a net qualitative difference between the behavior of the two cell lines: the aggregates of F98 cells contract and clearly organize in three dimensions, whereas the U87 cell aggregates keep an almost two-dimensional organization. So the superimposition parameter *α*_max_ should be larger for the F98 than for the U87 cells, see Fig 1 and 3. We can clearly detect when the parameter is too small, because then, the mean aggregate area in the simulations is too large, even for very small aggregates at the beginning of the experiment and even if the evolution of the number of aggregates is correct, see the red stars with *α*_max_ = 0.2 in S1 Fig (c). It is more difficult to detect when the parameter is too large: the difference between the cyan stars with *α*_max_ = 0.95 and the green stars with *α*_max_ = 0.7 in S1 Fig (c) is not obvious. Suppose that *α*_max_ = 1, all the cells in an aggregate could in theory superimpose and the area of any aggregate could be reduced to the area of a single cell. But since the motion step length of cells diminishes with the number of neighbors in aggregates, this process takes a lot of time, and it is not possible to see the complete superimposition of all the cells in an aggregate, during the time of experiments. We thus decided to infer the value of *α*_max_ from images of superimposition of two cells, see Fig 3 (b). From these images, it is clear that F98 cells allow a minimal distance between the cell centers smaller than the U87 cells, and we chose the value of *α*_max_ = 0.7 for the F98 cell line and *α*_max_ = 0.2 for the U87 cell line.

Two parameters still need to be set: the step length of individual cells and the proliferation rate. We determined the step length of individual cells so that the decrease of the number of aggregates corresponds to experimental data: if cells move too slowly, this number decreases also too slowly compared to experimental data (see Fig 6, cyan stars). If the step length is too large, the number of aggregates decreases too fast compared to experimental data (see Fig 6, red stars). The green stars correspond to the best value of the step.

**Fig 6.**
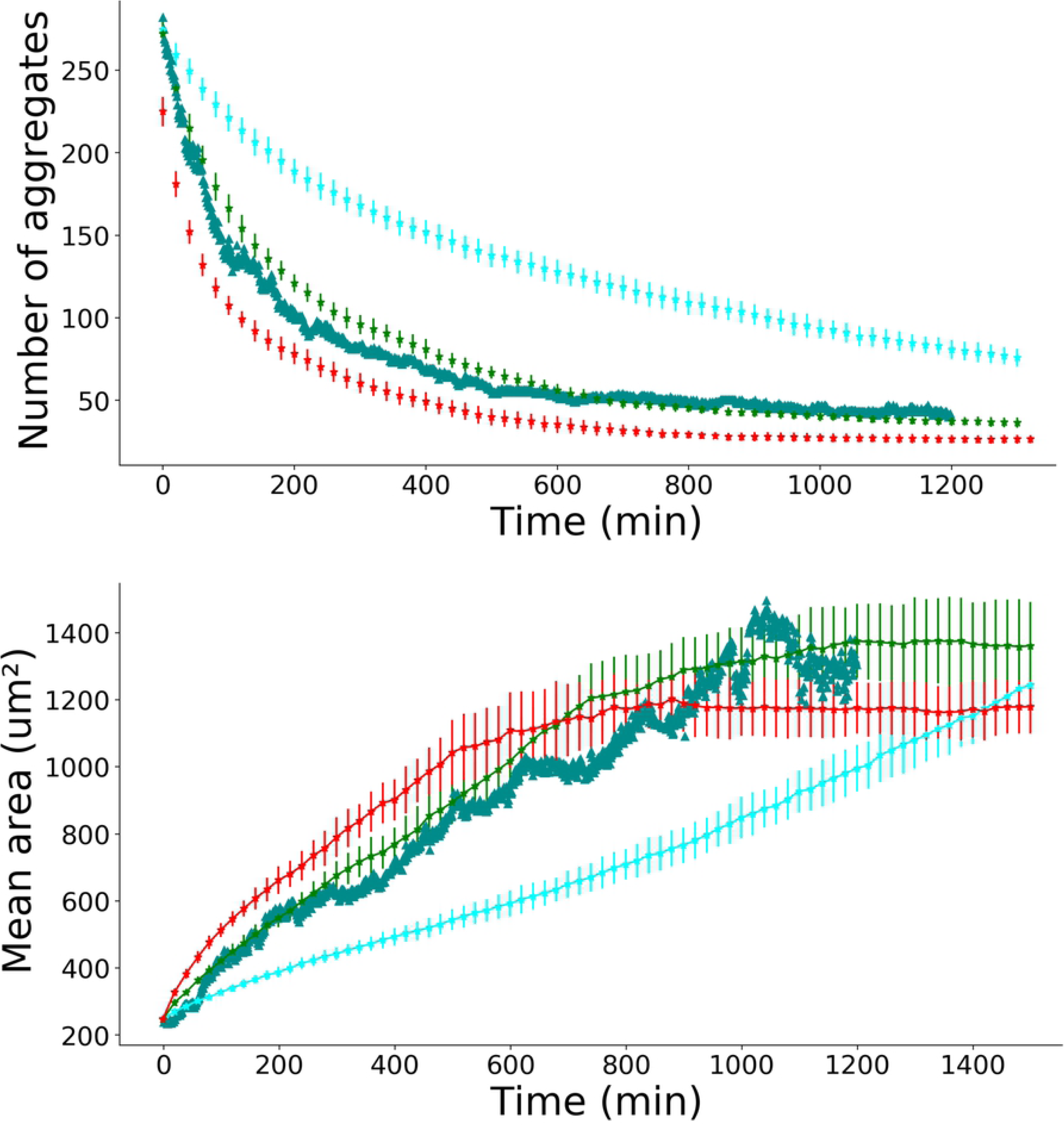
Parameter choice: step length. Experimental data points (one of the experiment on a non-adhesive substrate) are represented by green triangles (F98 cell line). Simulations: cyan stars, *a*_0_ = 1 pixel length; green stars, *a*_0_ = 6 pixel length; red stars, *a*_0_ = 14 pixel length. All other parameters are fixed: *κ* = 7 10^−4^ min^−1^, *α*_max_ = 0.7, *N*_0_ = 400, flux is on. The mean aggregate area evolution over time (bottom part) is using the same color scheme.

**Fig 7.**
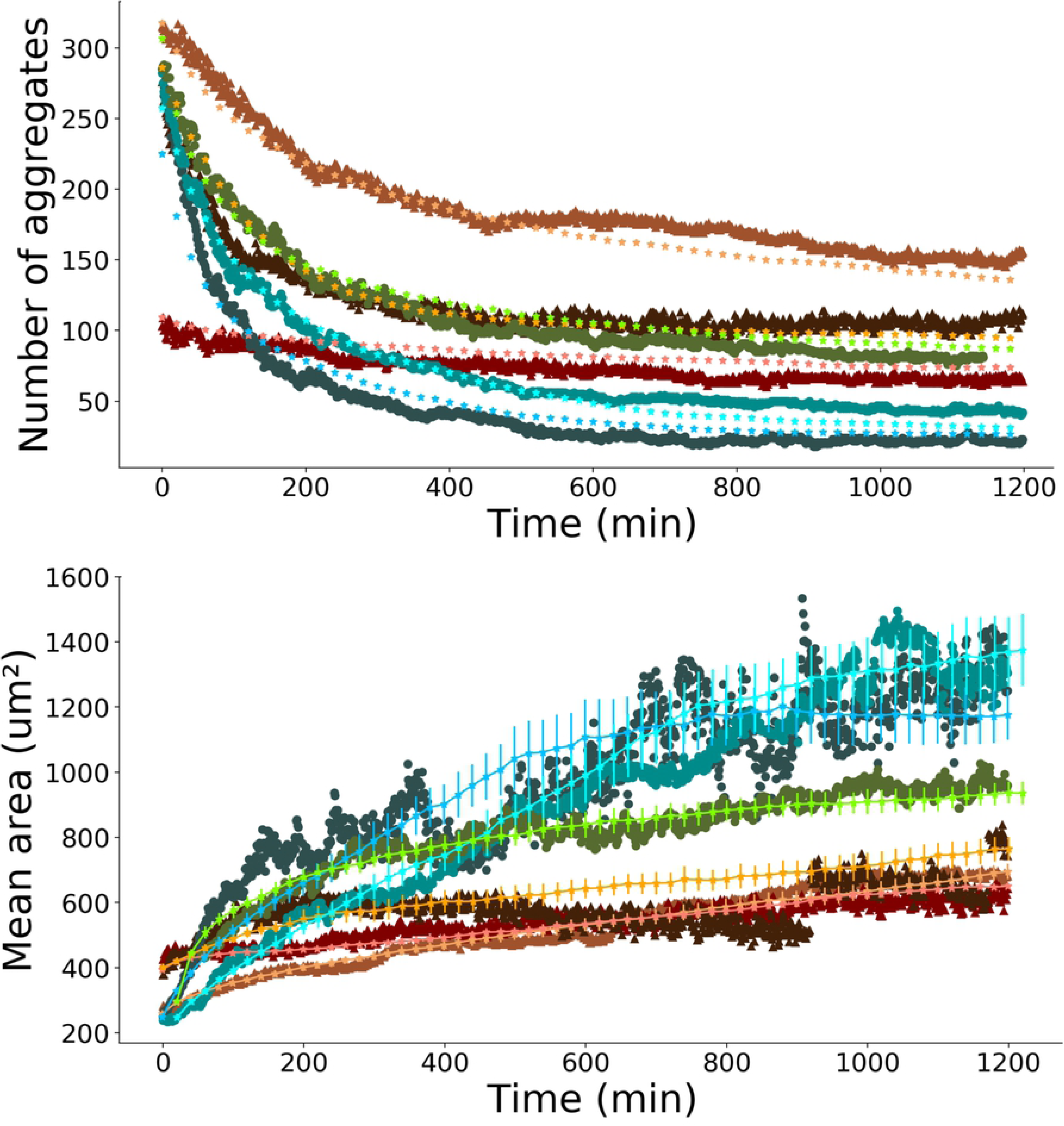
Experiments and simulations for the F98 cell line. Brown curves are for the adhesive case, blue/green ones are for the non-adhesive one. Triangles are for the experiment with adhesive substrate and circles are the non-adhesive substrate, while stars are for simulations. The upper part gives the evolution during time of the number of aggregates while the bottom part gives the mean area of aggregates over time. Simulation parameters: *a*_0_ = 3.7 pixel length, *N*_0_ = 400, flux is on (cyan); *a*_0_ = 6 pixel length, *N*_0_ = 784, flux is on (blue); *a*_0_ = 4.4 pixel length, *N*_0_ = 784, flux is on (green); *a*_0_ = 2 pixel length, *N*_0_ = 324, flux is off (orange); *a*_0_ = 1 pixel length, *N*_0_ = 781, flux is off (pink); *a*_0_ = 1.6 pixel length, *N*_0_ = 484, flux is off (brown). The parameters *α*_max_ and *κ* are common to all the simulations: *α*_max_ = 0.7, *κ* = 7 10^−4^ min^−1^. Error bars represent the standard deviation over 20 simulations.

The proliferation rate was chosen so that the mean area in simulations would fit the corresponding experiment at large times (the increase in the aggregate area after formation is only due to proliferation), see S1 Fig (b). The value *κ* = 10^−4^ min^−1^ (cyan stars) is too small, the value *κ* = 1.4 10^−3^ min^−1^ is too large and the value *κ* = 7 10^−4^ min^−1^ (green stars) is correct. We added the bias of the flux if visible in the experiments. Without any flux, aggregates still move but their motion is very small and the distance between them is too large to allow any collision. When the flux on individual as well as on aggregates is on, that corresponds to the green stars in S1 Fig (a), collisions are possible between large aggregates and the number of aggregates decreases even at large times.

We managed to reproduce the dynamics of both the mean area and the number of aggregate of each experiment and for both cell lines used. Results are shown in Fig 6 for the F98 cell line and in S2 Fig for the U87 cell line and for both adhesion conditions.

## 4 Discussion

We present here a combination of experimental and simulation results on the behavior of a cell population on soft hydrogels. On these soft gels, cells stay round, move and stick to each other to create aggregates. The shape and the size of these aggregates depend on the nature of the gels (adhesive or not adhesive), but also on the cell line: U87 aggregates are less cohesive than F98 aggregates, and for both cell lines, aggregates are smaller and more numerous on adhesive substrate (with PLL).

First, we compared the experimental data with the solutions of perikinetic equations. We found that the experimental non-adhesive and adhesive cases correspond to two different scenarii: in the non-adhesive dynamics of aggregation, a constant kernel leads to a better agreement with the experimental data, whereas the adhesive case is well fitted by a kernel that is non-zero only for particles of size 1. We found the kernel value *K* = 2.6 10^−13^ m^2^s^−1^ in the non-adhesive case, and a four-time smaller value of *K*_1_ in the adhesive case *K*_1_ = 6.4 10^−14^ m^2^s^−1^. These values are consistent with other studies [30], where only the case of a constant kernel is compared to experimetnal data. The agreement with the theoretical spaceless model is fair, but the model describes only the evolution of the number of aggregates as a function of time, whereas in our experiments, the area was also recorded.

We thus developed an agent-based model with simple rules that could reproduce as well the first stages of the experiments, when cells still move as individual cells, as the late stage where cells are in aggregates, and that could reproduce the experimental evolution of both the number and the mean area of the aggregates. To model all these stages, without describing precisely the shape of the cells, we estimated that a cellular Potts model was less adapted to our problem, compared to a classical agent-based cellular automaton. We introduced four rules: the motion rule (for individual cells, for cells inside an aggregate and for aggregates, in the presence or not of a flux), the superimposition rule, the proliferation rule and the compaction rule.

The superimposition and the compaction rules may need further justifications: Since the precise 3D organization in aggregates concerns only the F98 cell line, and to follow the approach of [13] where a stacking index is defined, we decided to keep our model in 2D and introduce an effective parameter of superimposition, that describes the strength of cell-cell adhesion and their ability to organize in 3D. This approach has the advantage of simplicity, since only one parameter can resume the difference between the behavior of the two cell lines.

The compaction that arises after the formation of cellular aggregates in general and is a collective effect of a cell population. In experiments, contraction of aggregates is due to individual cell contraction but also to the formation of supracellular stress cables, at the scale of the whole aggregate ([32]). This made us define a compaction rule that is non-local: the cells’ motion is biased towards the center of their aggregate and this bias increases with the size of the aggregate. Actually, the limitation of the motion is not severe: the maximum bias (in very big aggregates) restricts only the cell motion to a half plane towards the center of the aggregate.

With this agent-based model, in 2D and with simple rules, we were able to reproduce the behavior of two cell lines, namely the evolution of the number of aggregates and of their projected area, on two different substrates, one adhesive (with PLL) and one not (without PLL). More importantly, by fitting the number of aggregates and the mean area of aggregates as a function of time, we were able to infer quantitatively several properties of the two cells lines, on the two substrates: their speed of motion, their proliferation rate, their superimposition coefficient and the capacity of aggregates to compact.

First, our model allowed us to conclude that the effect of the presence of PLL in the gel (more adhesive substrate), for both cell lines, could be modeled as a simple slowing effect on cells. On non-adhesive hydrogels, there is often flux (probably due to inhomogeneities of the gels) which give to the cells and aggregates a motion bias (direction in only a half plane), that was taken into account in the model. We found that in the cellular automaton, in order to model the adhesive substrate, we had to decrease the step and remove the flux (i.e. remove the restriction of the motion to a half plane), for F98 cells, the mean speed motion of F98 is 4.7 ± 0.7 pixel length min^−1^=3.1 ± 0.4 *µ*m min^−1^ on non-adhesive substrates, whereas it is equal to 1.5 ± 0.3 pixel length min^−1^=1.0 ± 0.2 *µ*m min^−1^ on adhesive ones. For U87 cells the mean speed of motion are respectively 6 pixel lengths min^−1^=3.9 *µ*m min^−1^ and 2 pixel length min^−1^=1.3 *µ*m min^−1^.

The surface density of PLL molecules estimated from the volume concentration of 0.001% (w/v) PLL in the hydrogel precursor solution for a hydrogel thickness of 2 mm, is about 5 10^11^ molecules mm^−2^. About 5 10^5^ PLL molecules are found every *µ*m^2^. The F98 and U87 MG cells have a radius between 8 and 20 *µ*m so they move on a quasi-homogeneous surface of PLL molecules. The cells make smaller steps on PLL hydrogels because they are constrained in their motion by the electrostatic interactions they form with the PLL.

We also had to change the proliferation rate between the two cell lines. We found that the proliferation rate (in aggregates) is 7.10^−4^ min^−1^ for the F98 and 3.10^−4^ min^−1^ for the U87 cell line. Our results also reveal that the adhesive properties of the substrate does not impact the proliferation rate in aggregates: it is the same in the two conditions (adhesive and non-adhesive substrate), for each cell line.

The U87 cells are characterized by a weak adhesion between cells, leading to loose aggregates, whereas the F98 cells are much more cohesive. Moreover, there is no late compaction of the aggregates and the aggregates stay in 2D instead of organizing in 3D as in the F98 case. The results for both lines could be reproduced: in order to describe the U87 cell line we had to use a superimposition parameter smaller (*α*_max_ = 0.2) than the one used for the F98 line (*α*_max_ = 0.7) and to remove the compaction rule.

For two cell lines, we show here that by using our spatial model to fit the temporal evolution of the number of aggregates and their mean area, it is possible to infer the quantitative values of speed motion, proliferation and the qualitative abilities of cells to adhere to each other and to contract. It should be possible to study any cell line, providing that the gel stiffness is optimized for this cell line (indeed, carcinomas develop on a much stiffer substrate than gliomas, this was confirmed by a preliminary study of ours on the breast cancer cell line MCF7 which could not form aggregates on soft gels, dying rapidly). It has been shown for example that the cohesivity of aggregates (due to cell-cell adhesion) could be a clinically important parameter, since it seems to be inversely proportional to the *in vitro* invasive potential [33]. One promising direction of research is that of the study of cell lines from cancers which are known to develop metastases, such as breast cancer. In that case, our experimental technique should be adapted. Another interesting direction of future studies would be to modulate the adhesion. This could not be done in the present studies using PLL since a higher concentration of this molecule becomes toxic for the cells. Using an other adhesion molecule, such as RGD instead of PLL, we expect to be able to vary adhesion and study the aggregation phenomena for a wide range of values of the latter. We expect to return to address these problems, both experimentally and through modeling, in some future work of ours.

## Supporting information

**S1 Fig Parameter choice for F98 cell line.** Experimental data points (one of the experiment on a non-adhesive substrate) are represented by green triangles (F98 cell line). (a) Flux choice: no flux (cyan stars), flux on individual cell (red stars) and flux on individual cells and small aggregates (green stars). All other parameters are fixed: *a*_0_ = 6 pixel, *α*_max_ = 0.7, *κ* = 7 10^−4^ min^−1^, *N*_0_ = 400. Error bars are chosen equal to the standard deviation over 20 simulations. (b) Proliferation choice: cyan stars, *κ* = 10^−4^ min^−1^; green stars, *κ* = 7 10^−4^ min^−1^); red stars, *κ* = 1.4 10^−3^ min^−1^. All other parameters are fixed: *a*_0_ = 6 pixel length, *α*_max_ = 0.7, *N*_0_ = 400, flux is on. (c) Superimposition parameter choice: cyan stars, *α*_max_ = 0.95; green stars, *α*_max_ = 0.7; red stars, *α*_max_ = 0.2. All other parameters are fixed: *a*_0_ = 6 pixel length, *κ* = 7 10^−4^ min^−1^, *N*_0_ = 400, flux is on. In all the simulations, error bars represent the standard deviation over 20 simulations.

**S2 Fig Experiments and simulations for the U87 cell line.** Brown curves are the adhesive case, blue are for the non-adhesive case, triangles are for adhesive experiment and circles are for non adhesive experiment, stars are for simulations. The upper part gives the evolution during time of the number of aggregates while the bottom part gives the mean area of aggregates over time. Simulation parameters : *a*_0_ = 6 pixel length, *N*_0_ = 900, flux is off (cyan); *a*_0_ = 6 pixel length, *N*_0_ = 625, flux is off (blue); *a*_0_ = 2 pixel length, *N*_0_ = 484, flux is off (brown). The parameters *α*_max_ and *κ* are common to all the simulations: *α*_max_ = 0.2, *κ* = 3 10^−4^ min^−1^. Error bars represent the standard deviation over 20 simulations.

**S1 Video Aggregation of F98 cells** The aggregation process for the F98 cell line, during the first 400 minutes.

**S2 Video Aggregation of U87 cells** The aggregation process for the U87 cell line, during the first 600 minutes.

## Acknowledgments

This work was supported through a grant from Région Ile-de-France (‘DIM Problématiques transversales aux systèmes complexes ISC-2014-PME-003”) and a grant from the GEFLUC association (association des Entreprises Françaises dans la Lutte contre le Cancer).

